# Multinomial Convolutions for Joint Modeling of Sequence Motifs and Enhancer Activities

**DOI:** 10.1101/2020.07.28.224212

**Authors:** Minjun Park, Salvi Singh, Francisco Jose Grisanti Canozo, Md. Abul Hassan Samee

## Abstract

Massively parallel reporter assays (MPRAs) have enabled the study of transcriptional regulatory mechanisms at an unprecedented scale and with high quantitative resolution. However, this realm lacks models that can discover sequence-specific signals *de novo* from the data and integrate them in a mechanistic way. We present MuSeAM (<u>Mu</u>ltinomial CNNs for <u>Se</u>quence <u>A</u>ctivity <u>M</u>odeling), a convolutional neural network that overcomes this gap. MuSeAM utilizes multinomial convolutions that directly model sequence-specific motifs of protein-DNA binding. We demonstrate that MuSeAM fits MPRA data with high accuracy and generalizes over other tasks such as predicting chromatin accessibility and prioritizing potentially functional variants.

## Background

Proper regulation of gene transcription is critical to all forms of life. Regulatory sequences, also known as enhancers, are the key players in transcriptional regulation [1–4]. Enhancers coordinate with sequence-specific transcription factors (TFs) to ensure the transcriptional regulation of their target genes. In particular, enhancers harbor binding sites for TFs -- upon binding at these sites, TFs regulate the transcription of the enhancers’ target genes. Mutations in enhancer sequences can impact TF binding and result in transcriptional misregulation, which is a hallmark of many human diseases [5–9]. Consequently, a primary focus of regulatory genomics is to model genomic sequences for their enhancer activity, *i.e*., the extent to which a sequence can change the transcription level of a gene. Such a model can reveal transcriptional mechanisms, identify novel enhancers, and rank genomic variants based on their potential impact on transcriptional regulation.

Although there is a rich literature on enhancer activity models built from model organism data [10–15], modeling enhancer activity is still an elusive goal in human regulatory genomics. Due to the qualitative nature and low-throughput of classical reporter assays, we lack sufficient quantitative data on enhancer activities of human genome sequences. A recently developed assay known as the massively parallel reporter assay (MPRA) shows promise to overcome these challenges [16]. In a single experiment, an MPRA can quantify enhancer activities of thousands of candidate sequences. Briefly, in MPRA, a candidate sequence is first placed in the context of a reporter gene in a synthetic construct. The enhancer activity of the sequence is then quantified as the reporter gene’s normalized transcript count in that construct [16]. With this high throughput tool for quantitative data generation, recent studies have shifted their focus on building quantitative enhancer activity models from MPRA data.

The enhancer activity of a sequence is mostly driven by the TFs that bind to the sequence [4]. As such, an enhancer activity model needs to reveal at least two mechanistic information. First, the model should identify putative TF binding sites in a given sequence. Secondly, the model should infer the regulatory effects of the corresponding TFs and integrate them to fit the sequence’s enhancer activity. Enhancer activity models built from model organism data have shown remarkable success in capturing these mechanistic details [4, 10–12, 14, 15]. However, unlike the case for model organisms, the exact set of TFs active in a given cellular context and their sequence-specificities (motifs) are generally unknown in the case of human and higher-order vertebrates. Consequently, state of the art MPRA models have used a broad collection of sequence-specific features. These features include motifs from motif databases, sequence features modeled from functional genomic datasets, sequence *k*-mers, and evolutionary conservation, GC-content, and DNA shape [17]. To improve model performance, current models have also utilized epigenetic features, such as histone tail modifications [17]. Since most of these features are not interpretable as specific TF binding sites, unfortunately the models lose their mechanistic interpretation. Overall, in the realm of enhancer activity modeling from MPRA data, it is an open question whether one can achieve both mechanistic interpretation and a satisfactory level of model performance.

We posit that a systematic, data-driven approach to jointly learn sequence-specific motifs and model enhancer activity from MPRA data can achieve satisfactory modeling performance without losing mechanistic interpretation. Here we propose MuSeAM (<u>Mu</u>ltinomial CNNs for <u>Se</u>quence <u>A</u>ctivity <u>M</u>odeling) -- a convolutional neural network (CNN) model that uses “multinomial convolutions” to achieve this goal. The use of CNNs in sequence data modeling is a well-established trend in genomics [18, 19]. However, interpreting both the convolutions and the deep architectures has remained difficult (Discussion). Here we show that a simple transformation can help us learn convolutions directly as multinomial distributions over the four nucleotides. Such multinomial distributions are directly interpretable as the classical motif models of TF-DNA binding specificity that have been utilized for decades in genomic sequence analyses. Searching public motif databases for matches to these multinomial convolutions, one can identify the corresponding TFs. Convolving a sequence with these multinomial convolutions gives us likelihood terms, which we use in a linear regression model to fit the sequence’s enhancer activity data. The coefficients in this linear regression model represent the regulatory effects and strength of the TFs. MuSeAM’s implementation is customizable, so one can replace the linear regression model with any other architecture of their choice.

As proof of concept, we applied MuSeAM on a recent, high quality MPRA dataset of more than 2000 human genomic sequences assayed in the HepG2 (human liver) cell-line (Inoue et al. 2019). Despite its simple formulation, MuSeAM achieved state of the art performance in modeling this data, while also learning multinomial convolutions that are directly interpretable as TF motifs. MuSeAM motifs (the multinomial convolutions) recapitulated the known TFs, their regulatory roles, and their co-occurrence patterns. Furthermore, MuSeAM learned a highly generalizable sequence model, as we demonstrate through the model’s ability in predicting genome-wide DNA accessibility, although accessibility data was not used in training the model. MuSeAM also prioritized common single-nucleotide polymorphisms (SNPs) for their potential to alter enhancer activity. The top SNPs are enriched for low minor allele frequencies and they mostly occur within promoters and enhancers in the human genome, highlighting MuSeAM’s ability to learn biologically meaningful models. Overall, MuSeAM was able to retain high mechanistic interpretability along with viable performance, a feat not realized by other state of the art models. We therefore believe that MuSeAM’s multinomial convolution approach will have a significant impact on future studies to build interpretable neural network models of genomic sequence data.

## Results

### MuSeAM: A Multinomial Convolutional Neural Network for Sequence Activity Modeling

MuSeAM (<u>Mu</u>ltinomial CNNs for <u>Se</u>quence <u>A</u>ctivity <u>M</u>odeling) is a Convolutional Neural Network model of enhancer activity that offers clear interpretability through a new class of convolutions in the convolutional layer and a simple linear function in the fully connected layer.

#### Multinomial convolutions

Convolution matrices in our model satisfy the conditions of multinomial distributions over the four DNA nucleotides. In particular, given a convolution matrix *X*, we transform *X* into a matrix *T* such that:

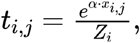

where:

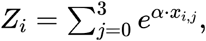

and *α* is a free parameter of the model (Fig. 1A, Method). Here *i* and *j* are indices for the rows and the columns of *T*, respectively.

**Figure 1.**
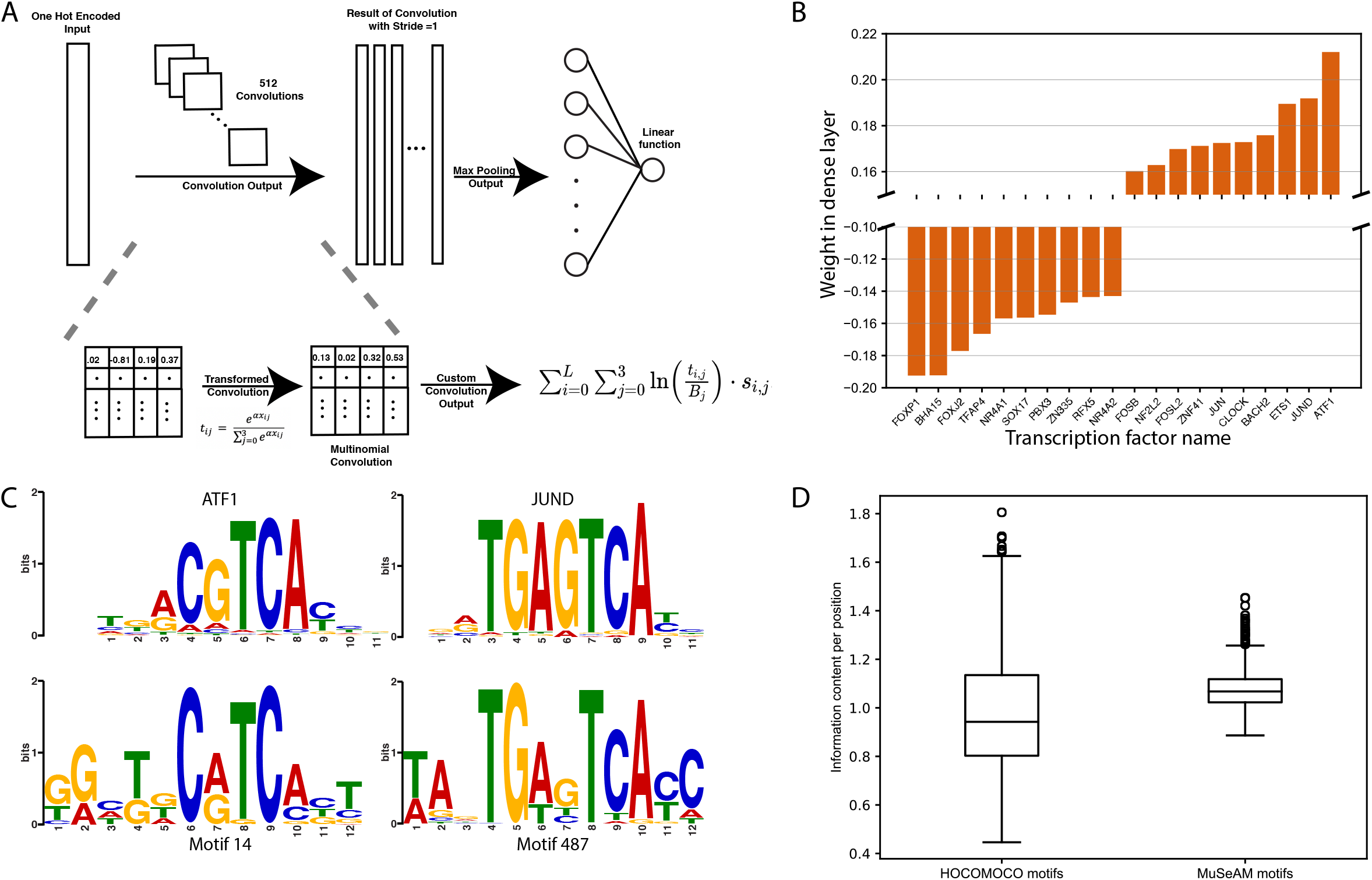
(A) Overview of the MuSeAM model, (B) The ten strongest activators and repressors, along with their weights in the fully connected layer (the linear regression function), as identified by the MuSeAM model, (C) Comparisons between known motifs (top) and MuSeAM motifs (bottom) for top two activators, (D) Information content per position for known motifs (left) and MuSeAM motifs (right).

Thus, each row of the matrix *T* satisfies the conditions of a multinomial distribution:

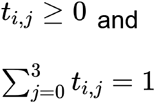

These multinomial convolutions are directly interpretable as the widely used motif models of TF-DNA binding specificity. In the remainder of the paper, we use the term *MuSeAM motifs* to denote the multinomial convolutions of a trained MuSeAM model.

Given a multinomial convolution matrix *T* of size *L* × 4 and a “one-hot encoded” (Methods) nucleotide sequence *S*, the convolution operation in MuSeAM computes the following value at each *L*-length subsequence *s* of *S*:

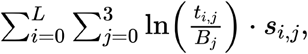

where *B* is a background distribution over the four DNA nucleotides. We call this value the LLR (logarithm of likelihood ratio) term for *s*, since it denotes the logarithm of the ratio of likelihoods for generating *s* from *T* versus from *B*. We pass the LLR terms through a ReLU activation and a max-pooling function; the output of the max-pooling layer is the motif score for *T* in *S* [14, 20]. Thus, for a multinomial convolution *T* and a sequence *S*, we consider only the best matching subsequence *s* for *T* in *S*. However, if every *s* in *S* is a better match to the background distribution, then we do not consider any subsequence from *S* (motif score is zero).

#### Linear regression function

The fully connected layer in our model implements a linear function combining the motif scores, resembling two other classical models of enhancer activity [14, 20]. Thus, from a mechanistic perspective, MuSeAM implements the bag-of-binding sites or billboard model [21]. The coefficients of this linear model then represent the strength and the role (activating or repressing) of each TF.

### MuSeAM Models MPRA Data in the Human HepG2 Cells

To demonstrate the utility of multinomial CNNs in learning regulatory motifs and modeling enhancer activity, we trained MuSeAM on Inoue *et al*.’s recent MPRA dataset [22]. In this dataset, Inoue *et al*. reported the enhancer activity of 2440 sequences assayed in the HepG2 cell-line. They synthesized each sequence with 100 unique barcodes to obtain 100 independent enhancer activity values for the sequence, making the estimated enhancer activities highly robust. Moreover, a recent study has reported a detailed model of this dataset (see below), which provides a means to benchmark our model.

In a 10-fold cross-validation setting, the optimal MuSeAM model fit Inoue *et al*.’s dataset with a coefficient of determination of 0.43 and a Spearman correlation coefficient of 0.54 (mean over ten folds; standard deviations: 0.058 and 0.067, respectively) (Supplemental Table S1). This optimal model used 512 convolutions, each modeling TF-DNA binding specificity over 12 bps (convolution size 12 × 4). This performance is on a par with the Spearman correlation coefficient of 0.59 ± 0.04 reported by Kreimer *et al*. in their recent extensive modeling of this dataset [17]. In this study, Kreimer *et al*. utilized five models (Extra Trees, Gradient Boosting, Random Forest, Elastic Net, and an ensemble model that averages the first four models) on more than 5500 features. The features included predictions from other deep learning models (DeepBind [19] and DeepSEA [18]), motifs from motif databases, k-mers, epigenetic marks, and general sequence features such as GC content, length of polyA/T stretch, DNA shape features, and evolutionary conservation. Because of the exhaustive nature of Kreimer *et al*’s study, we consider their performances as practical limits for a model on this dataset, and it was remarkable that despite its simplicity and clear interpretability, MuSeAM performed on a par with Kreimer *et al*’s results.

### MuSeAM Reveals Relevant Transcription Factors and Their Regulatory Roles

To understand whether MuSeAM has learned biologically relevant motif models, we performed a series of analyses. We first ensured that the MuSeAM motifs are comparably informative to experimentally derived motifs. We compared the information content per position (ICP) of the MuSeAM motifs against the ICP of motifs from motif databases (Fig. 1B). In fact, this revealed that the ICP values of MuSeAM motifs are higher than the known motifs (Welch two sample t-test, p-value < 2.2e-16; Methods).

Next we checked if MuSeAM motifs correspond to TFs with known regulatory function in the human liver. It is a fundamental expectation that an MPRA model will identify the TFs that regulate the reporter gene’s expression [23]. Such a model should also infer each TF’s role, *i.e*., whether it is an activator or a repressor of gene expression. It is straightforward in MuSeAM to find both information. First, searching motif databases [24, 25] for matches to each MuSeAM motif identifies the relevant TF. Secondly, the fully connected layer weight corresponding to the max pooled value of a MuSeAM motif provides the information on the TF’s role and strength. In particular, the sign of this weight denotes whether the corresponding TF is an activator (positive weight) or repressor (negative weight), and the magnitude of the weight suggests how strong the TF is in this role. Of note, since MuSeAM fits the multinomial convolutions in a data-driven manner, a database search may not necessarily find a hit for every multinomial convolution. These examples would be indicative of novel motifs.

Following the above rationale, we took the MuSeAM motifs corresponding to the ten strongest activators and repressors from the trained model (Fig. 1B; Supplemental Table S2). A motif database search for these MuSeAM motifs found matching TFs with similar regulatory roles as implicated by the model (Fig. 1C; Supplemental Table S2; Methods).

For example, MuSeAM identified ATF1 (Activating Transcription Factor 1) as the strongest activator for this MPRA dataset. ATF1 is a member of the CREB family of cAMP-responsive activators in mammalian systems. The TF is expressed ubiquitously [26] and is known for its role as a transcriptional activator [27–29]. Similarly, MuSeAM identified FOXP1 (Forkhead Box P1) as the strongest repressor in the MPRA dataset. FOXP1 is expressed in the liver [30] and the TF acts as a repressor of transcription [31, 32]

We also found a high concordance between the most “influential” TFs and the strongest regulators (activator/repressor) in this model. We define the influence score of a TF as the average increase in MuSeAM’s error on this dataset if we drop the corresponding MuSeAM motif from the model. Of the top 10% most influential TFs, about 80% are among the strongest activators and repressors. For example, the five most influential TFs according to this analysis are FOXP1, ETS1, FOXJ2, BACH2, and TFAP4. From these, ETS1 and BACH2 are among the five strongest activators, and FOXP1, FOXJ2, and TFAP4 are among the five strongest repressors.

Finally, we checked if MuSeAM motif pairs have captured any co-binding patterns known for regulatory TFs. Of note, since we formulated the model as a “bag of sites” model and the max-pooling operation only considers the strongest matches to the convolutions, we expected that the number of MuSeAM motif pairs recapitulating known co-binding patterns would be rather small. Still, in our analysis of MuSeAM motif pairs for enriched co-occurrence within 12 bps (a distance threshold characteristic to TF complexes [33] (Methods), we found 1108 (~21%) MuSeAM motif pairs indeed capture the formation of known TF complexes. The top ten TF pairs having the highest enrichment in this analysis are FOSL2-JUND, HNF1B-HNF1A, FOSL2-FOS, JUND-FOS, BACH2-FOSL2, ATF1-FOSL2, FOSL1-JUND, FOS-FOSL1, ATF2-FOSL2 and ATF1-ATF2. Interestingly, all of these pairs are known for protein-protein interaction in BioGRID [34] or STRING [35].

Overall, these analyses suggested that MuSeAM has indeed learned biologically relevant motif models, encouraging us to utilize the model to elicit more biological information.

### MuSeAM Discovers Distinct Groups of Regulatory Sequences in Terms of Motif Composition

Studies in model organisms suggest that enhancers can be clustered into groups according to the TFs that target the different enhancer groups. Identifying these enhancer clusters are important in revealing the transcriptional regulatory mechanisms in a cell-type. Since Inoue *et al*. selected their candidate sequences based on enhancer marks (ChIP-seq peaks of EP300 and H3K27ac), we posited that these sequences would cluster into smaller groups based on the most important TFs for each sequence.

We defined a TF’s importance for a sequence as the increase in error in predicting the sequence’s enhancer activity if we remove the MuSeAM motif corresponding to that TF from the model. Upon clustering the sequences according to their ten most important TFs, we found seven clusters (Fig. 2A; Methods). A differential analysis between the clusters identified the characteristic TFs for each cluster (Fig. 2B; Supplementary Figure S1; Methods). For example, the most important TFs characterizing cluster 3 are FOSL1, BACH2, HNF1B, HNF1A, CEBPE, and FOSL2. In general, each cluster has at least one TF that uniquely characterizes it. For example: JUND characterizes cluster 0 and SP3 characterizes cluster 1. Interestingly, this analysis separated the 200 synthetic control sequences (Cluster 3, Fig 2A: Methods) in Inoue *et al*’s dataset from the real genomic sequences, further underscoring that MuSeAM has learned meaningful biological signals from this dataset.

**Figure 2.**
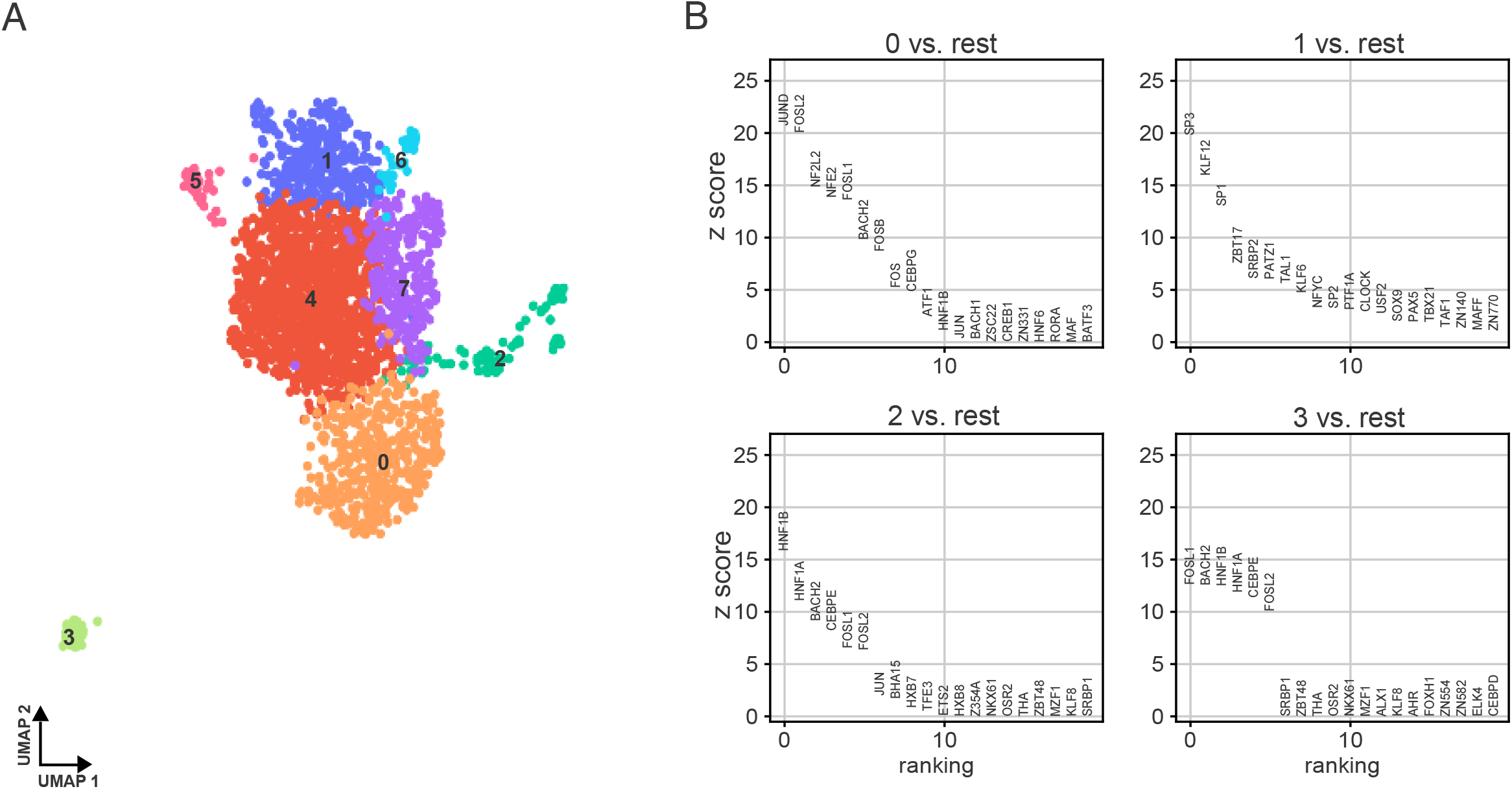
(A) Clustering the sequences in the dataset based on the ten most important transcription factors (TFs) for each sequence, as identified by the MuSeAM model. Clusters were computed through uniform manifold approximation and projection (UMAP) followed by Leiden clustering, and (B) Cluster-specific TFs identified by a Wilcoxon rank-sum test comparing each of the first four clusters against all other clusters. Characteristic TFs for the clusters: Cluster 0 – JUND, Cluster 1 – SP3, Cluster 2 – HNF1B, Cluster 3 – FOSL1.

### MuSeAM Suggests a Sequence Specific Explanation of Accessibility from DNase-Seq Data

Since MuSeAM learns and integrates sequence-specific signals to model an MPRA dataset, we reasoned that a trained MuSeAM model could explain other functional annotations that depend on the same sequence-specific signals. Previous studies have demonstrated that accessibility of genomic DNA is a sequence-specific feature largely explained by TF motifs, and accessibility of a sequence is strongly associated with its enhancer activity [36–39]. Thus, we sought to see if the trained MuSeAM model can explain genomic accessibility.

For this analysis, we used the ENCODE consortium’s [40] uniformly processed DNase-Seq peaks from the HepG2 cell-line (approximately 200K peaks). We applied MuSeAM to predict the enhancer activities of these peak sequences. The predicted enhancer activities showed a remarkable agreement with the accessibility levels across DNase-Seq peaks, ranked and grouped according to their accessibility. In particular, we found a Pearson correlation coefficient of −0.88 between the rank of a group and the median of predicted enhancer activity within the group (Fig. 3A; Methods).

**Figure 3.**
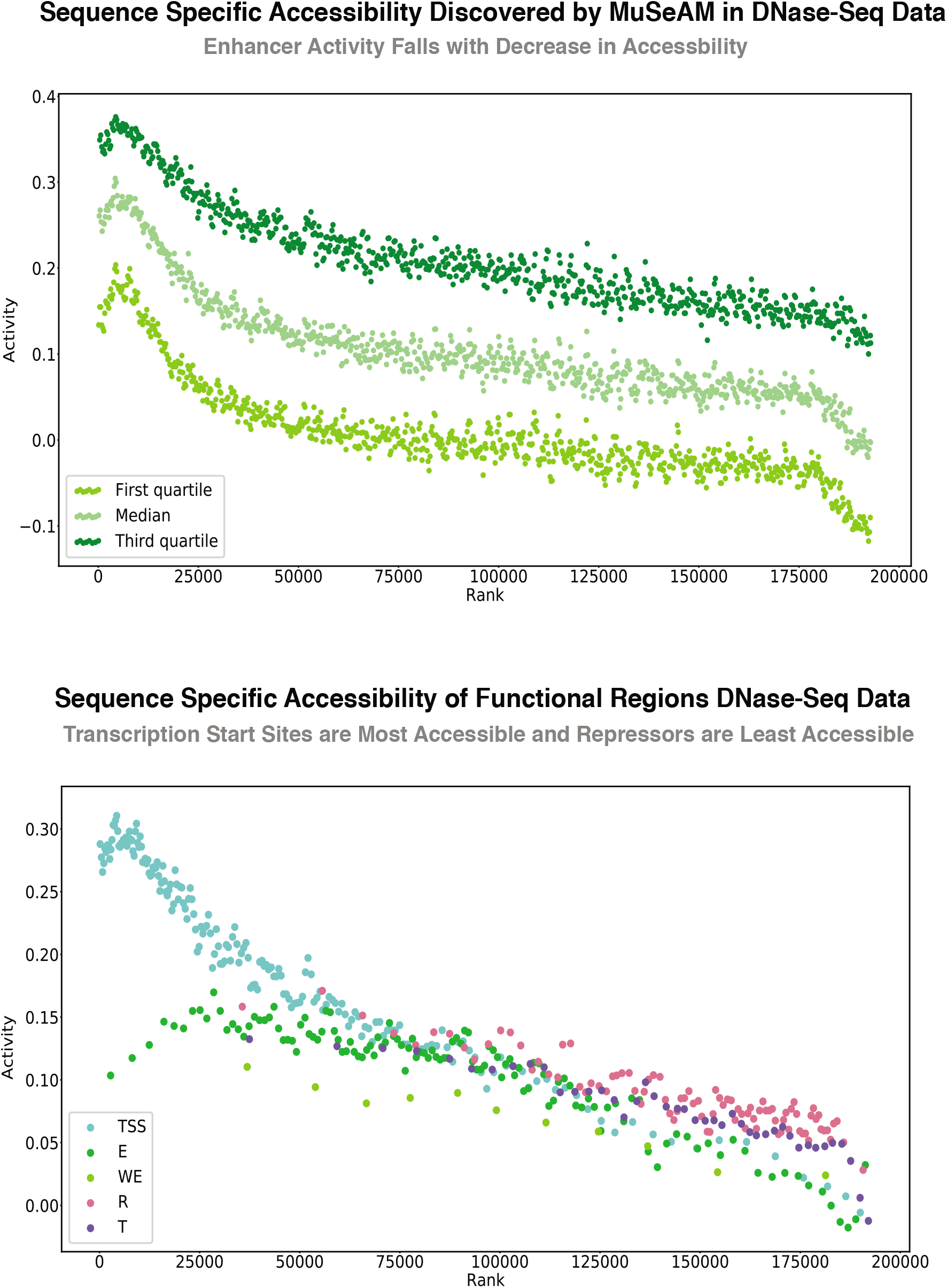
(A) MuSeAM models genome-wide chromatin accessibility for DNase-Seq peaks in the HepG2 cell-line, accurately predicting decrease in activity with decreasing accessibility, (B) For DNase-Seq peaks in HepG2 cell-line annotated as enhancers and transcription start sites, MuSeAM predicts high levels of activity, whereas the repressed regions correspond to low activity level predictions. TSS: Transcription Start Sites, E: Enhancers, WE: Weak Enhancers, R: Repressors, T: Transcription Sites

A comparison against the functional annotations (such as enhancers, transcription start sites, repressed regions, etc.) of genomic sequences in the HepG2 cell-line [40] further suggested that the trained MuSeAM model has learned sequence-specific features that characterize different functional regions in the genome. We found that MuSeAM predicted the highest levels of enhancer activities for DNase-Seq peaks that are transcription start sites and enhancers, while the predicted activities dropped consistently for the peaks that represent repressed regions (Fig. 3B; Methods).

Since the above analyses did not require retraining the MuSeAM model on DNase-Seq data, it suggests that the model is capable of learning sequence-specific signals that characterize a broad range of regulatory elements.

### MuSeAM Identifies Functionally Important Variants among Common SNPs

Given MuSeAM’s success in modeling enhancer activity from MPRA and in predicting genome-wide accessibility signals, we wanted to utilize the model further to predict the potential effect of genomic variants. We assumed that the change in the enhancer activity of a sequence because of a genomic variant, as predicted by MuSeAM, is a surrogate for its functional impact.

For this analysis, we focused on the common bi-allelic variants from dbSNP [41], *i.e*., variants having a minor allele frequency (MAF) of at least 1%, that overlap with DNase-Seq peaks in the HepG2 cell-line (Methods). For each SNP, we took the 151 bp long genomic sequence centered at the SNP, and applied MuSeAM to compute the enhancer activities of the major and the minor allele versions of this sequence. We defined the SNP’s *predicted variant effect* (PVE) as the difference between these two values of enhancer activity (Methods). The predicted variant effects for the 131K common variants showed a skewed pattern, with a median of 0.014 and range of 0.0--0.441 (Fig. 4A; Methods).

**Figure 4.**
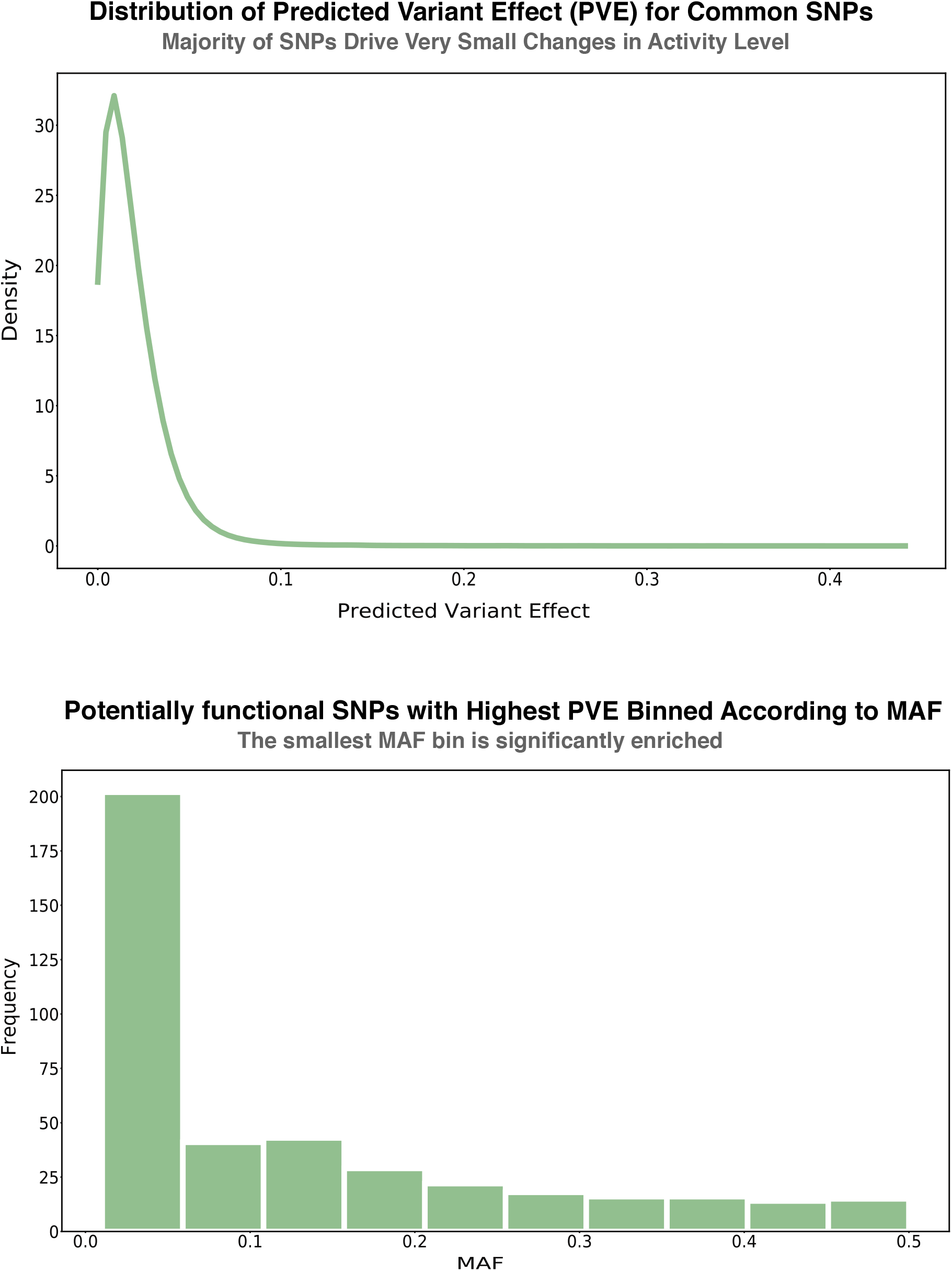
(A) Distribution of variant-driven difference in activity (Predicted Variant Effect or PVE) as predicted by MuSeAM, for dbSNP’s common variants within DNase peaks in HepG2 cell-line, (B) Distribution of potentially functional SNPs according to their Minor Allele Frequencies

To get a conservative list of variants that are potentially functional, *e.g*., are able to alter enhancer activity or genomic accessibility, we then selected the top 1% common SNPs according to their PVE values and applied an FDR correction based on the PVEs of benign SNPs. At 5% FDR, we found 416 common SNPs. These SNPs are significantly enriched for low minor allele frequency (MAF) (Fig 4B; Methods). Although there is no benchmark study to make a conclusive statement about the functional relevance of these SNPs, our findings aligned with the hypothesis that a variant whose presence is a strong driver of change in activity, is likely to have lower minor allele frequencies (MAF) [42–44]. Furthermore, these potentially functional SNPs disrupt genomic regions annotated for activity as enhancers, with 62% of the SNPs in the smallest MAF bin overlapping with enhancers or promoters. At these genomic regions, the SNPs disrupt potential binding sites for TFs like BACH2, NF2L2, NFE2, and FOSL2, that are known to play important roles in the human liver and were also found as important regulators by MuSeAM for the MPRA dataset.

## Methods

### MPRA Data

We collected Inoue *et al*.’s [22] MPRA dataset of 2,440 sequences and their enhancer activities. Each sequence was chromosomally integrated in the HepG2 cell-line with as many as 100 barcodes. The large number of barcodes ensured reproducible and quantitative measurements of regulatory activity, and controlled against nonuniformity in oligonucleotide synthesis [22].

### Motif Databases

To compare the MuSeAM motifs against currently known motifs, we used motifs from the HOmo sapiens COmprehensive MOdel COllection (HOCOMOCO)-v11 [24] database of transcription factor binding models. The current HOCOMOCO release includes 769 motifs representing the DNA-binding specificity of 680 TFs.

### Accessibility Data

We collected the ENCODE consortium’s [40] uniformly processed DNA accessibility data (DNaseI hypersensitivity peaks) for the HepG2 cell-line (identifier: ENCSR000EJV) from: http://hgdownload.soe.ucsc.edu/goldenPath/hg19/encodeDCC/wgEncodeAwgDnaseUniform/wgEncodeAwgDnaseUwdukeHepg2UniPk.narrowPeak.gz.

### Genome Segmentations According to Functional Groups

We collected the ENCODE consoritum’s [40] genome segmentation data (from DNase, FAIRE, Histone and TFBS Signals) for the HepG2 cell-line from: http://hgdownload.soe.ucsc.edu/goldenPath/hg19/encodeDCC/wgEncodeAwgSegmentation/wgEncodeAwgSegmentationCombinedHuvec.bed.gz

### Common SNP Data

We collected the common single nucleotide polymorphisms from dbSNP [41] (build 151) from https://hgdownload.soe.ucsc.edu/goldenPath/hg19/database/snp151Common.txt.gz.

### Protein-Protein Interaction Data

We searched for protein-protein interactions in the STRING [35] and the BioGRID [34] databases.

### MuSeAM: A Multinomial Convolutional Neural Network Model

MuSeAM is a convolutional neural network model with a custom-implemented convolutional layer and a fully connected layer. Specifically, the model architecture comprises the input layer that takes DNA sequences, the convolutional layer, a maximum pooling layer, and finally, a dense layer whose output is the predicted expression level for the input sequence. The model was implemented using Keras in TensorFlow [45].

#### Input Layer

For each sequence, the model receives two sets of one-hot encoded inputs: one is a representation of the forward strand of the sequence, and the other is the reverse complement of the same. The one-hot encoding of a genomic sequence *S* of length *N* is an *N* × 4 matrix Enc(*S*) where the *i*-th row is a vector encoding the identity of the i-th nucleotide of *S* as follows.

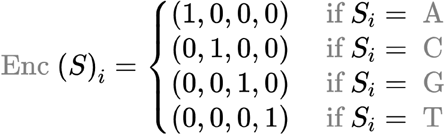

For the ease of notation, we write *S* to denote both the DNA sequence and its one-hot encoding.

#### Convolution Layer

##### Multinomial Convolution Matrices

The convolution layer in MuSeAM is a custom implementation where each convolution learns multinomial distributions. In particular, given a convolution matrix *X*, we transform *X* into a matrix *T* such that:

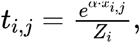

where:

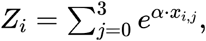

and *α* is a free parameter learned by the model. Here *i* and *j* are indices for the rows and the columns of *T*, respectively.

Thus, each row of the matrix *T* satisfies the conditions of a multinomial distribution:

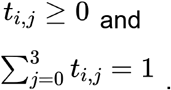

Note that the convolutions in our model do not use any trainable bias term.

#### Convolution and pooling operation

For each multinomial convolution matrix *T* of size *L ×* 4 and a one-hot encoded nucleotide sequence *S* of length *N* as described previously, a convolution operation computes the following term on each *L*-length subsequence *s* of *S*.

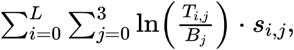

where *B* is a background distribution over the four DNA nucleotides. In other words, for each *L*-length subsequence *s* of *S*, the convolution operation computes the logarithm of the ratio of two likelihood terms. The first likelihood (the numerator) is the likelihood for generating the sequence *s* from the convolution matrix *T* (*i.e*., its multinomial distribution). The second likelihood is for generating *s* from the background. In our current analysis, we used the uniform distribution as our background distribution *B*.

The convolution operation produces a vector of *N* – *L* + 1 log-likelihood ratio (LLR) terms for each of the two incoming input units (corresponding to the forward-strand and the reverse-complement representations of the input sequence *S*). We pass the LLR terms through a ReLU activation and a max-pooling function; the output of the max-pooling layer is the motif score for *T* in *S* [14, 20]. Thus, for a multinomial convolution *T* and a sequence *S*, we consider only the best matching subsequence *s* for *T* in *S*. However, if every *s* in *S* is a better match to the background distribution, then we do not consider any subsequence from *S* (motif score is zero).

Note that the transformations to construct the multinomial distributions do not change the original convolution matrices. As such, we do not need to customize how the parameter optimization algorithm updates the convolution weights.

#### Fully Connected Layer

For each sequence *S*, the motif scores representing the strongest LLR of each convolution *T* are flattened and passed to the final dense layer. This layer implements a linear regression function (linear activation function) with L1 regularization to fit the expression value for *S*.

### Optimization and Model Selection

#### Loss Function

MuSeAM optimises the parameters using the mean squared error (MSE) function as the loss function. Given a batch of *n* sequences with the true and the predicted values of the i-th sequences being *y_i_* and *ŷ_i_*, respectively, the MSE function computes the following term.

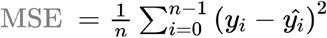

In our implementation, we used batches of size 512.

#### Model Optimization

We compiled MuSeAM to optimize MSE loss with the *adam* optimizer and reported the model performance in terms of coefficient of determination and Spearman’s rank-order correlation.

All weights and bias terms in the model are randomly initialized with the *gloret uniform* initializer.

#### Model Selection

We used 10-folds cross validation to select the optimal values for the following parameters: number of convolutions, size of each convolution, regularization type, number of epochs, and batch size. For each fold, we used 10% data for validation. We recorded the average coefficient of determination and Spearman correlation coefficient across the ten folds and used mean coefficient of determination to determine the best set of learning parameters.

### Searching for MuSeAM Motif Matches in Other Motif Databases

We used the tool tomtom from the MEME Suite [46] to search for MuSeAM motif matches in the HOCOMOCO database [24].

### Statistical Test Comparing Information Content Per Position of HOCOMOCO and MuSeAM Motifs

Given a motif *M* of length *L*, we first normalize the elements in each row of *M* by the sum of the row. Then we calculated the information content per position (ICP) of the normalized motif as:

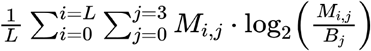

Here *B* represents the background distribution. In the current analysis, we used the uniform distribution. Given the ICP values of the HOCOMOCO motifs [47] and the MuSeAM motifs, we performed a Welch two-sample t-test using the R command `t. test (datal, data2, alternative=“less”, var. equal=False)`, where datal and data2 are the ICP values of HOCOMOCO and MuSeAM motifs, respectively.

### Statistical Test for Enriched Co-occurrence of MuSeAM Motifs

For each pair of MuSeAM motifs, we first count the number of sequences where they co-occur. We say that a MuSeAM motif *occurs* in a given sequence if the motif’s score (*i.e*., its maximum LLR score) is positive in that sequence. Next we count the number of sequences where the pair of MuSeAM motifs co-occur within 12 bps. We chose the threshold of 12 bps since the DNA binding sites for TFs in a complex are almost always located within this distance [33]. From these two values, we then computed a Binomial p-value to test whether co-occurrences of the pair of MuSeAM motifs are enriched within the distance threshold. After correcting for multiple hypotheses tests, at a p-value threshold of 10^−3^, we find 1109 MuSeAM motif pairs, corresponding to 73 TFs, to have enriched co-occurrences within 12 bps distance.

### Computing the Influence Score of MuSeAM Motifs

Let *ŷ_i_* denote MuSeAM’s prediction for the *i*-th sequence in the MPRA dataset. Let *ŷ_i,j_* denote MuSeAM’s prediction for the *i*-th sequence upon removing the *j*-th convolution from the model. We define the influence score of the *j*-th convolution (MuSeAM motif) as:

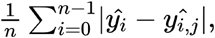

where *n* denotes the number of sequences in the dataset.

### Clustering Sequences According to Important MuSeAM Motifs

Similar to the idea of an importance score of a MuSeAM motif with respect to the entire dataset, we also defined the motif’s importance score per sequence as follows. Let *ŷ_i_* denote MuSeAM’s prediction for the *i*-th sequence. Let *ŷ_i,j_* denote MuSeAM’s prediction for the *i*-th sequence upon removing the *j*-th convolution from the model. We define the importance score of the *j*-th convolution (MuSeAM motif) for the *i*-th sequence as: |*ŷ_i_* – *ŷ_i,j_*|

We chose for each sequence the ten most important MuSeAM motifs. We identified all the unique transcription factors (183 in total) represented by these MuSeAM motifs, and created a matrix with the TFs as columns and the sequences as rows. A cell in this matrix was assigned a value of 1 if the corresponding TF was important for the corresponding sequence, otherwise we assigned 0 to the cell. This matrix was then used as input to identify any meaningful clustering of sequences that might be implicit based on the MuSeAM motifs.

Using principal component analysis (PCA) [48], we first computed a low dimensional manifold that best explains the variance in the matrix above, and further reduced the dimensionality into two by using the top 50 principal components in a uniform manifold approximation and projection (UMAP) [49]. We then clustered the sequences in this two dimensional space using the Leiden community detection algorithm [50]. We used this pipeline from the Scanpy package [51].

To detect TFs that are significantly over-represented in each cluster as compared to all other clusters, we performed a Wilcoxon rank-sum test [52]. We corrected the resulting p-values at an FDR threshold of 5% by using the Benjamini-Hochberg procedure [53].

### Application of MuSeAM on DNase-Seq Data

For this analysis, we used the ENCODE consortium’s [40] DNA accessibility data for the HepG2 cell-line (identifier: ENCSR000EJV). It comprises 192,959 accessible sequences (DNaseI hypersensitivity peaks) along with a measure of their accessibility. The majority of peak sequences in this dataset are 150 bps in length. For the remaining peaks, we considered only the central 150 bp sequence in our analysis.

For the plots shown in Fig. 3, we sorted the peaks according to their accessibility and grouped them in sliding windows of size 500 (shift size 250) along this sorted list. Within each window, we applied MuSeAM to predict the enhancer activities for all sequences in that window and we recorded the first three quartiles of this distribution.

To annotate a DNase-Seq peak according to its potential function, we used the ENCODE consortium’s [40] genome segmentations (from DNase, FAIRE, Histone and TFBS Signals) for the HepG2 cell-line. We annotated a peak with a segmentation label (such as enhancer, transcription start site, etc.) if at least half of the peak overlaps with a genomic region of that label. For the annotated peaks, we used MuSeAM’s predicted activity level, and plotted the median activity with median rank for each category of annotations to examine the relationships if any between the accessibility based on the functional annotations and predicted activity (Fig 3B).

### Identifying Functionally Important Variants from Common SNPs

From dbSNP’s dataset of common SNPs (build 151), we considered the bi-allelic SNPs that overlap with DNase-Seq peaks in the HepG2 cell-line. For each of the 131,405 SNPs selected above, we considered the 151 bp sequence centered at the SNP and created two sequences that differ only at the central position by having either the major or the minor allele. The difference in MuSeAM predicted enhancer activities of these two sequences is the predicted variant effect (PVE) statistic for the SNP.

For the top 1% (largest PVE) SNPs from this analysis (1304 SNPs), we applied a false discovery rate (FDR) correction using the Benjamini-Hochberg procedure [53]. To this end, we collected the list of benign SNPs from dbSNP [41] and created a null distribution of PVE (search string: `(“benign”[Clinical Significance]) AND “snv”[SNP Class])`. At a 5% FDR, we obtained 416 SNPs that we considered for further analysis.

To test whether SNPs with strong PVE are enriched for low minor allele frequencies (MAFs), we binned the SNPs into ten equal intervals according to their MAF values. Under the null hypothesis that SNPs are distributed uniformly across the bins, but we found the first bin to be significantly enriched for strong PVE SNPs (Binomial test p-value less than 1e-50).

The SNPs in the bin with the lowest MAF values were mapped to functional annotations (such as enhancers, transcription start sites, repressed regions, etc.) collected from the ENCODE [40] consortium.

## Discussion

Massively parallel reporter assays (MPRAs) hold great promise in eliciting transcriptional regulatory mechanisms in a biological system of interest [16, 54, 55]. Modeling MPRA datasets can identify the putative transcription factors (TFs) acting on the assayed sequences, the regulatory roles of these TFs and their relative influence on transcriptional regulation. However, because of the lack of appropriate models, the goal of answering these mechanistic questions is still elusive. State of the art models achieve high accuracy but lack the interpretability that is necessary to answer these mechanistic questions.

Here we proposed MuSeAM (<u>Mu</u>ltinomial CNNs for <u>Se</u>quence <u>A</u>ctivity <u>M</u>odeling), a multinomial convolutional neural network (multinomial CNN) for modeling MPRA dataset. We found that MuSeAM is able to answer all the mechanistic questions mentioned above while achieving high performance. Importantly, utilizing the powerful feature discovery techniques of convolutional neural networks, MuSeAM answers the mechanistic questions in a data-driven way; the model does not require any prior knowledge of the particular biological system.

The key to MuSeAM’s success is the idea of multinomial convolutions which, to our knowledge, has not been discussed before in the literature. The use of CNNs is becoming increasingly common for modeling genomic datasets [18, 19, 56, 57]. These models are particularly useful for discovering sequence-specific signals in the input sequences and integrating the strength of these signals to model a quantitative value for the sequences. However, interpretability of CNN models has remained a concern [58]. In fact, it is not expected that a CNN will readily highlight interpretable relationships in data or will guide the formulation of mechanistic hypotheses [59].

The multinomial convolution formulation is particularly attractive because of its immediate interpretability as the widely used sequence-motifs, without making any assumption about the concentration of TFs [57] or requiring any post-hoc analysis [58]. Furthermore, the resulting MuSeAM motifs are interoperable with the large array of genomic data analysis pipelines that are currently utilized for applications like sequence annotation and motif enrichment. We therefore anticipate that multinomial convolutions will find broader use in the future. One can always utilize the multinomial formulation as the first convolution layer with an arbitrarily complex downstream architecture. As another appealing extension, we envision more biophysically grounded functions [60] to replace our pooling and fully connected layer.

In the current application, MuSeAM not only answered the main mechanistic questions on transcriptional regulation from Inoue *et al*.’s dataset (Inoue et al. 2019), but it also showed remarkable accuracy in predicting the levels of genome-wide chromatin accessibility in the same cell-line. The model also short-listed SNPs that carry independent evidence of being functional, implying that the model indeed is capable of discovering and generalizing sequence-specific signals from the data. Of note, a GWAS variant’s effect on gene expression is often modest [61]. Therefore, while we expect the SNPs with large PVEs to be functionally important, we anticipate that our conservative approach has missed many functional variants.

Given MuSeAM’s success in achieving state of the art accuracy, it was interesting for us to ask *“when does the model fail?”*. In a follow-up analysis to characterize the “problematic” sequences, *i.e*., the sequences for which MuSeAM consistently made erroneous predictions, we found that these sequences have extremely high enhancer activity (Supplemental Figure S2A). In particular, for each fold during the 10-fold cross validations, we sorted the test sequences according to MuSeAM’s error in predicting their enhancer activity and selected the top 5% sequences as the problematic sequences. To see if problematic sequences appear across models with different numbers of convolutions, we used Jaccard Index (JI). We found a mean JI of 0.881 (range 0.846 -- 0.92), suggesting that it is the same set of sequences, *i.e*., sequences with very high enhancer activity, on which MuSeAM consistently failed. Although MuSeAM models with more multinomial convolutions generally fit these sequences better, using more than 512 convolutions did not show major improvement in the mean coefficient of determination for the overall dataset (Supplemental Figure S2B).

Finally, since this is a central assumption that the model will converge at convolutions that are interpretable as the conventional motifs, it is interesting to ask “*how does a multinomial convolution learn specificity signals?”* We indeed found that the MuSeAM motifs have an average information content per position that is generally higher than the known motifs. We note that, the gradient of the output of the convolution operation with respect to its input, *i.e*., 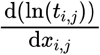, reduces to 1 – *t_i,j_*. In other words, minimizing the loss function is equivalent to maximizing the *t_i,j_* terms, which represent probabilities in the multinomial distributions. Thus, under the formulation of multinomial convolution, the optimization process ensures learning specificity signals while also fitting the data.

## Supplementary Figure Legends

**Figure S1.**
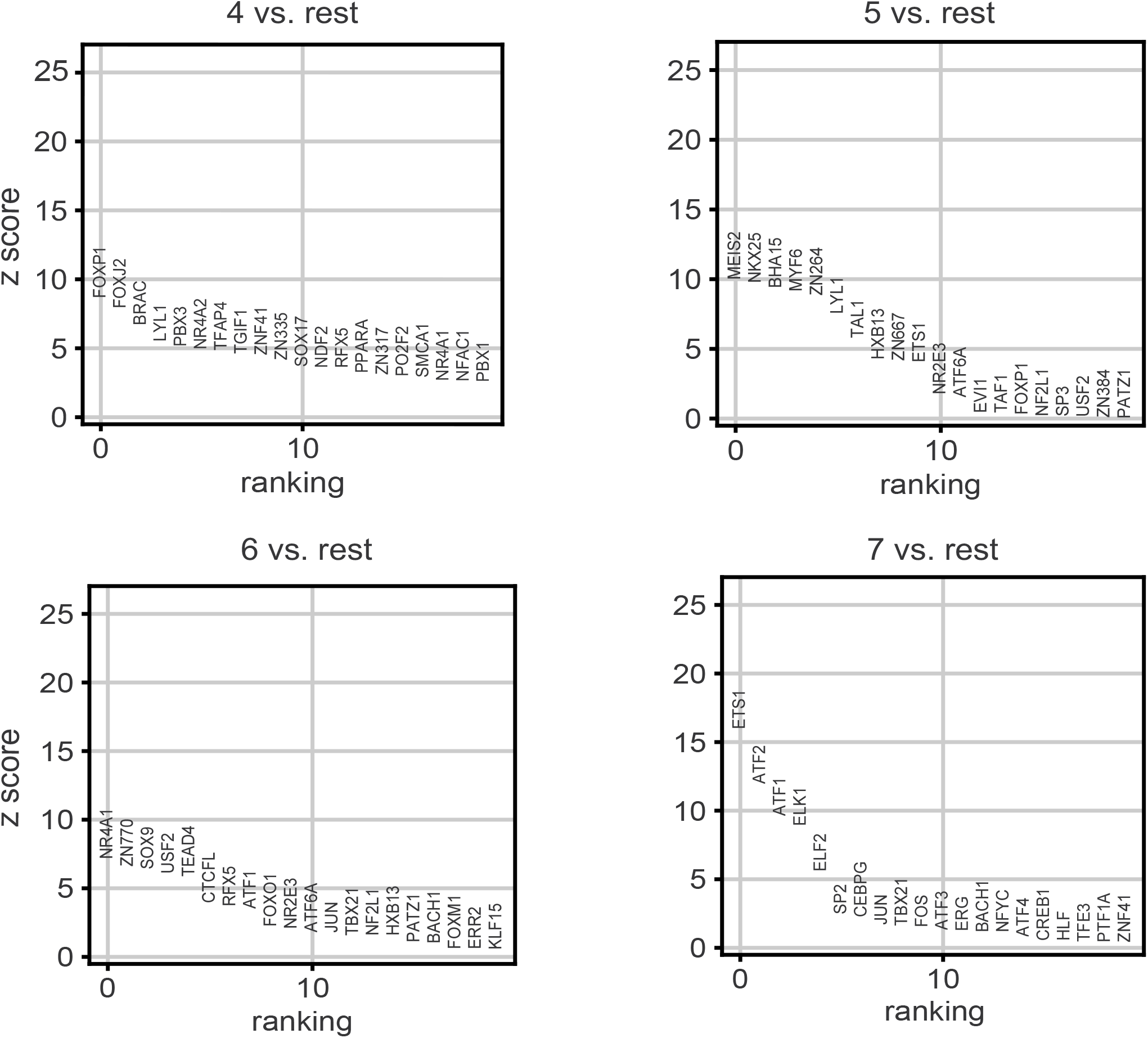
Wilcoxon rank-sum test of one versus all for the clusters 4-7. Characteristic TFs for the clusters: Cluster 4-FOXP1, Cluster 5 – MEIS2, Cluster 6 – NR4A1, Cluster 7 – ETS1.

**Figure S2.**
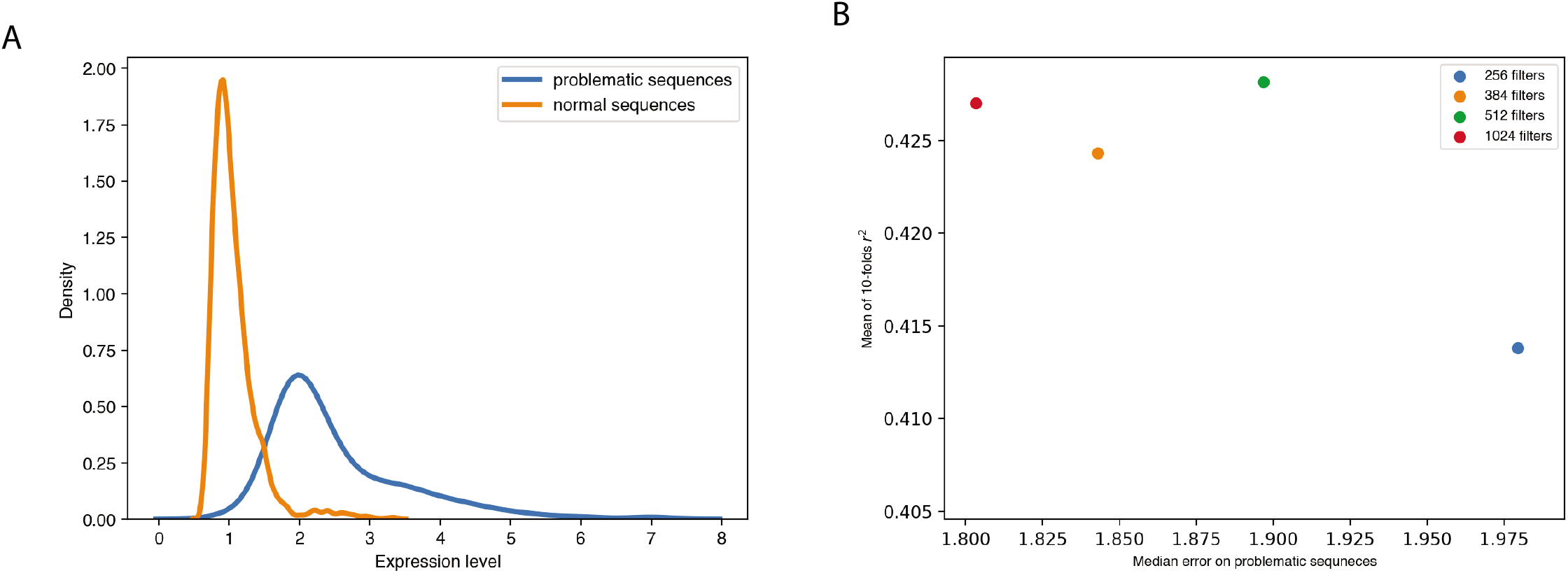
(A) Density plot of enhancer activity levels for all sequences (orange) and problematic sequences (blue). (B) With an increase in the number of convolutions, the model performs better on the problematic sequences, but does not show major improvement in overall performance. Shown are the mean coefficient of determination values for all sequences vs. median error for problematic sequences.

## Supplementary Table Titles

Table S1. Summary of MuSeAM’s modeling performance over 10 folds with coefficient of determination, Pearson coefficient correlation and Spearman’s rank correlation coefficient as reporting metrics.

Table S2. Summary of strongest activator and repressor motifs learnt by MuSeAM along with corresponding TFs from HOCOMOCO motif database.

## References

1. Levo M, Segal E (2014) In pursuit of design principles of regulatory sequences. Nat Rev Genet 15:453–468

2. Shlyueva D, Stampfel G, Stark A (2014) Transcriptional enhancers: from properties to genome-wide predictions. Nat Rev Genet 15:272–286

3. Weingarten-Gabbay S, Segal E (2014) The grammar of transcriptional regulation. Hum Genet 133:701–711

4. Spitz F, Furlong EEM (2012) Transcription factors: from enhancer binding to developmental control. Nat Rev Genet 13:613–626

5. Cherry TJ, Yang MG, Harmin DA, et al (2020) Mapping the cis-regulatory architecture of the human retina reveals noncoding genetic variation in disease. Proc Natl Acad Sci U S A 117:9001–9012

6. Xu D, Gokcumen O, Khurana E (2020) Loss-of-function tolerance of enhancers in the human genome. PLoS Genet 16:e1008663

7. Chatterjee S, Ahituv N (2017) Gene Regulatory Elements, Major Drivers of Human Disease. Annu Rev Genomics Hum Genet 18:45–63

8. Lee TI, Young RA (2013) Transcriptional regulation and its misregulation in disease. Cell 152:1237–1251

9. Farooq M, Troelsen JT, Boyd M, et al (2010) Preaxial polydactyly/triphalangeal thumb is associated with changed transcription factor-binding affinity in a family with a novel point mutation in the long-range cis-regulatory element ZRS. Eur J Hum Genet 18:733–736

10. Fakhouri WD, Ay A, Sayal R, et al (2010) Deciphering a transcriptional regulatory code: modeling short-range repression in the Drosophila embryo. Mol Syst Biol 6:341

11. Gertz J, Siggia ED, Cohen BA (2009) Analysis of combinatorial cis-regulation in synthetic and genomic promoters. Nature 457:215–218

12. Segal E, Raveh-Sadka T, Schroeder M, et al (2008) Predicting expression patterns from regulatory sequence in Drosophila segmentation. Nature 451:535–540

13. Bintu L, Buchler NE, Garcia HG, et al (2005) Transcriptional regulation by the numbers: models. Curr Opin Genet Dev 15:116–124

14. Bussemaker HJ, Li H, Siggia ED (2001) Regulatory element detection using correlation with expression. Nat Genet 27:167–171

15. Beer MA, Tavazoie S (2004) Predicting gene expression from sequence. Cell 117:185–198

16. Inoue F, Ahituv N (2015) Decoding enhancers using massively parallel reporter assays. Genomics 106:159–164

17. Kreimer A, Yan Z, Ahituv N, Yosef N (2019) Meta-analysis of massively parallel reporter assays enables prediction of regulatory function across cell types. Human Mutation 40:1299–1313

18. Zhou J, Troyanskaya OG (2015) Predicting effects of noncoding variants with deep learning-based sequence model. Nat Methods 12:931–934

19. Alipanahi B, Delong A, Weirauch MT, Frey BJ (2015) Predicting the sequence specificities of DNA- and RNA-binding proteins by deep learning. Nature Biotechnology 33:831–838

20. Kazemian M, Blatti C, Richards A, et al (2010) Quantitative analysis of the Drosophila segmentation regulatory network using pattern generating potentials. PLoS Biol 8.: https://doi.org/10.1371/journal.pbio.1000456

21. Arnosti DN, Kulkarni MM (2005) Transcriptional enhancers: Intelligent enhanceosomes or flexible billboards? J Cell Biochem 94:890–898

22. Inoue F, Kircher M, Martin B, et al (2017) A systematic comparison reveals substantial differences in chromosomal versus episomal encoding of enhancer activity. Genome Res 27:38–52

23. Ernst J, Melnikov A, Zhang X, et al (2016) Genome-scale high-resolution mapping of activating and repressive nucleotides in regulatory regions. Nat Biotechnol 34:1180–1190

24. Kulakovskiy IV, Vorontsov IE, Yevshin IS, et al (2018) HOCOMOCO: towards a complete collection of transcription factor binding models for human and mouse via large-scale ChIP-Seq analysis. Nucleic Acids Res 46:D252–D259

25. Fornes O, Castro-Mondragon JA, Khan A, et al (2020) JASPAR 2020: update of the open-access database of transcription factor binding profiles. Nucleic Acids Res 48:D87–D92

26. Mayr B, Montminy M (2001) Transcriptional regulation by the phosphorylation-dependent factor CREB. Nat Rev Mol Cell Biol 2:599–609

27. Atlas E, Stramwasser M, Mueller CR (2001) A CREB site in the BRCA1 proximal promoter acts as a constitutive transcriptional element. Oncogene 20:7110–7114

28. Kingsley-Kallesen ML, Kelly D, Rizzino A (1999) Transcriptional Regulation of the Transforming Growth Factor-β2 Promoter by cAMP-responsive Element-binding Protein (CREB) and Activating Transcription Factor-1 (ATF-1) Is Modulated by Protein Kinases and the Coactivators p300 and CREB-binding Protein. Journal of Biological Chemistry 274:34020–34028

29. Lu Z, Sack MN (2008) ATF-1 is a hypoxia-responsive transcriptional activator of skeletal muscle mitochondrial-uncoupling protein 3. J Biol Chem 283:23410–23418

30. Uhlén M (2015) Tissue-based map of the human proteome. Science 347.: https://doi.org/10.1126/science.1260419

31. Sollis E, Graham SA, Vino A, et al (2016) Identification and functional characterization of de novo FOXP1 variants provides novel insights into the etiology of neurodevelopmental disorder. Hum Mol Genet 25:546–557

32. Jepsen K, Gleiberman AS, Shi C, et al (2008) Cooperative regulation in development by SMRT and FOXP1. Genes Dev 22:740–745

33. Jolma A, Yin Y, Nitta KR, et al (2015) DNA-dependent formation of transcription factor pairs alters their binding specificity. Nature 527:384–388

34. Stark C, Breitkreutz B-J, Reguly T, et al (2006) BioGRID: a general repository for interaction datasets. Nucleic Acids Res 34:D535–9

35. Szklarczyk D, Gable AL, Lyon D, et al (2019) STRING v11: protein-protein association networks with increased coverage, supporting functional discovery in genome-wide experimental datasets. Nucleic Acids Res 47:D607–D613

36. Catarino RR, Stark A (2018) Assessing sufficiency and necessity of enhancer activities for gene expression and the mechanisms of transcription activation. Genes Dev 32:202–223

37. Heinz S, Romanoski CE, Benner C, Glass CK (2015) The selection and function of cell type-specific enhancers. Nat Rev Mol Cell Biol 16:144–154

38. Thurman RE, Rynes E, Humbert R, et al (2012) The accessible chromatin landscape of the human genome. Nature 489:75–82

39. Hashimoto T, Sherwood RI, Kang DD, et al (2016) A synergistic DNA logic predicts genome-wide chromatin accessibility. Genome Res 26:1430–1440

40. ENCODE Project Consortium (2012) An integrated encyclopedia of DNA elements in the human genome. Nature 489:57–74

41. Kitts A, Phan L, Ward M, Holmes JB (2014) The Database of Short Genetic Variation (dbSNP). National Center for Biotechnology Information (US)

42. Montgomery SB, Lappalainen T, Gutierrez-Arcelus M, Dermitzakis ET (2011) Rare and common regulatory variation in population-scale sequenced human genomes. PLoS Genet 7:e1002144

43. Hernandez RD, Uricchio LH, Hartman K, et al (2019) Ultra-rare variants drive substantial cis-heritability of human gene expression. Nat Genet 51:1349

44. Zhao J, Akinsanmi I, Arafat D, et al (2016) A Burden of Rare Variants Associated with Extremes of Gene Expression in Human Peripheral Blood. Am J Hum Genet 98:299–309

45. Abadi M, Agarwal A, Barham P, et al (2016) TensorFlow: Large-Scale Machine Learning on Heterogeneous Distributed Systems. arXiv [cs.DC]

46. Bailey TL, Johnson J, Grant CE, Noble WS (2015) The MEME Suite. Nucleic Acids Res 43:W39–49

47. Kulakovskiy IV, Vorontsov IE, Yevshin IS, et al (2016) HOCOMOCO: expansion and enhancement of the collection of transcription factor binding sites models. Nucleic Acids Res 44:D116–25

48. Jolliffe I (2014) Principal Component Analysis. Wiley StatsRef: Statistics Reference Online

49. McInnes L, Healy J, Saul N, Großberger L (2018) UMAP: Uniform Manifold Approximation and Projection. Journal of Open Source Software 3:861

50. Traag VA, Waltman L, van Eck NJ (2019) From Louvain to Leiden: guaranteeing well-connected communities. Sci Rep 9:5233

51. Wolf FA, Angerer P, Theis FJ (2018) SCANPY: large-scale single-cell gene expression data analysis. Genome Biol 19:15

52. Gooch JW (2011) Wilcoxon Rank-Sum Test. Encyclopedic Dictionary of Polymers 1002–1002

53. Haynes W (2013) Benjamini–Hochberg Method. Encyclopedia of Systems Biology 78–78

54. Smith RP, Taher L, Patwardhan RP, et al (2013) Massively parallel decoding of mammalian regulatory sequences supports a flexible organizational model. Nat Genet 45:1021–1028

55. Gordon MG, Inoue F, Martin B, et al (2020) lentiMPRA and MPRAflow for high-throughput functional characterization of gene regulatory elements. Nat Protoc. https://doi.org/10.1038/s41596-020-0333-5

56. Kelley DR, Reshef YA, Bileschi M, et al (2018) Sequential regulatory activity prediction across chromosomes with convolutional neural networks. Genome Res 28:739–750

57. de Boer CG, Vaishnav ED, Sadeh R, et al (2020) Deciphering eukaryotic gene-regulatory logic with 100 million random promoters. Nat Biotechnol 38:56–65

58. Shrikumar A, Greenside P, Kundaje A (2017) Learning Important Features Through Propagating Activation Differences. In: International Conference on Machine Learning. pp 3145–3153

59. Eraslan G, Avsec Ž, Gagneur J, Theis FJ (2019) Deep learning: new computational modelling techniques for genomics. Nat Rev Genet 20:389–403

60. He X, Samee MAH, Blatti C, Sinha S (2010) Thermodynamics-based models of transcriptional regulation by enhancers: the roles of synergistic activation, cooperative binding and short-range repression. PLoS Comput Biol 6.: https://doi.org/10.1371/journal.pcbi.1000935

61. Gallagher MD, Chen-Plotkin AS (2018) The Post-GWAS Era: From Association to Function. Am J Hum Genet 102:717–730

